# A dialysis medium refreshment cell culture set-up for an osteoblast-osteoclast coculture

**DOI:** 10.1101/2022.09.15.508113

**Authors:** M.A.M. Vis, B.W.M. de Wildt, K. Ito, S. Hofmann

**Affiliations:** Orthopaedic Biomechanics, Department of Biomedical Engineering and Institute for Complex Molecular Systems (ICMS) Eindhoven University of Technology, Eindhoven, Netherlands

**Keywords:** Dialysis, Osteoblast, Osteoclast, Bone, Coculture

## Abstract

Culture medium exchange leads to loss of valuable auto- and paracrine factors produced by the cells. However, frequent renewal of culture medium is necessary for nutrient supply and to prevent waste product accumulation. Thus it remains the gold standard in cell culture applications. The use of dialysis as a medium refreshment method could provide a solution as low molecular weight molecules such as nutrients and waste products could easily be exchanged, while high molecular weight components such as growth factors, used in cell interactions, could be maintained in the cell culture compartment. This study investigates a dialysis culture approach for an *in vitro* bone remodeling model. In this model, both the differentiation of human mesenchymal stromal cells (MSCs) into osteoblasts and monocytes (MCs) into osteoclasts is studied. A custom-made simple dialysis culture system with a commercially available cellulose dialysis insert was developed. The data reported here revealed increased osteoblastic and osteoclastic activity in the dialysis groups compared to the standard non-dialysis groups, mainly shown by significantly higher alkaline phosphatase (ALP) and tartrate-resistant acid phosphatase (TRAP) activity, respectively. This simple culture system has the potential to create a more efficient microenvironment allowing for cell interactions via secreted factors in mono- and cocultures and could be applied for many other tissues.

## 1. Introduction

In conventional cell and tissue culture strategies, cells/tissues are surrounded by culture medium containing various nutrients and stimulants. This culture medium is exchanged regularly to maintain nutrient and stimulant levels upon cell consumption and to eliminate waste products produced by the cells [1]. While in culture, the cells also actively produce and secrete a variety of macromolecules to maintain their microenvironment. By changing the medium, these important molecules for auto- and paracrine signaling are lost. After every medium change the cells must make a new effort to restore their communication through these molecules. This effort could considerably influence the cell’s behavior. However, frequent renewal of the medium is necessary to provide sufficient nutrients and prevent waste product accumulation, which has been shown to inhibit cell growth [2].

To overcome this waste accumulation problem, cell culture systems have been developed in which culture medium was filtered or dialyzed to remove waste products [3] [4]. The principle of dialysis relies on the exclusion of molecules based on their size [1]. Fresh medium contains low molecular weight (MW) molecules such as nutrients, amino acids, and vitamins. Depending on the chosen MW cut-off (MWCO), the dialysis membrane allows for exchange of those molecules. In this way waste products can diffuse out of the culture medium while nutrients and vitamins can diffuse back in. High MW components such as growth factors produced by the cells or added to the medium are retained in the medium compartment [1].

This dialysis culture principle has already been reported in 1958 by G.G Rose who introduced the concept of using a cellophane membrane to divide the tissue culture part from the culture medium exchange part [3] [4]. He kept advancing the set-up and eventually engineered a 12-chambered tissue culture system with dialysis membranes [5]. Decades later, E.A. Vogler reported a cell culture device based on Rose’s work and showed long-term cell culture (30 days) of a variety of mammalian cells (epithelioid (canine), fibroblastic (monkey), hybridoma, primary thymocytes and splenocytes (all murine)) in the stable culture environment created by dialysis [6]. Vogler’s group further expanded this idea of “simultaneous growth and dialysis” and over the years reported work in the field of bone tissue engineering and breast cancer metastasis with cell cultures ranging from two-dimensional (2D) to three-dimensional (3D) [7] [8] [9] [10].

Currently, dialysis for re-use of culture medium is not frequently reported. It is remarkable that these techniques are not used more often in cell culture systems while they have the potential to improve cell function tremendously. Dialysis cultures have both economic and biological benefits. The economic benefits come from the fact that less of the costly macromolecules for cell proliferation and differentiation are needed in the culture medium. The biological benefit originates from the maintenance of the cells’ secretome over time, which leads to a more stable cell culture environment. This seems particularly imperative for studies investigating cell-cell-communication via cell-secreted macromolecules or extracellular vesicles. Through the retention of those factors in the medium, dialysis cultures could be beneficial for the proliferation and differentiation process of many cell types, particularly for the culture of *in vitro* tissue models. These models aim at representing the *in vivo* situation as closely as possible in order to for example test the effect of drugs on cells. It is hypothesized that a microenvironment providing more stability through less fluctuations could contribute to the efficacy and reliability of such models.

In the most recent study of Krishnan *et al*., Vogler’s bioreactor was used to create a 3D bone remodeling model with murine osteoblasts and osteoclasts cultured over a period of 2-10 months [10]. With a goal towards *in vitro* bone-remodeling models for drug testing and personalized medicine, a human model would be preferred over a murine model due to interspecies differences. Moreover, a 2-10 month culture period would generate practical issues for high throughput drug testing. Therefore, we propose a fully human 3D bone-remodeling model, including the use of human platelet lysate (hPL) instead of the generally used fetal bovine serum (FBS) [11]. We present a simple, cost-effective method for incorporating dialysis into the cell culture. A silk fibroin (SF) scaffold will be integrated to reduce the amount of time needed to create the 3D bone tissue (Figure 1A).

**Figure 1.**
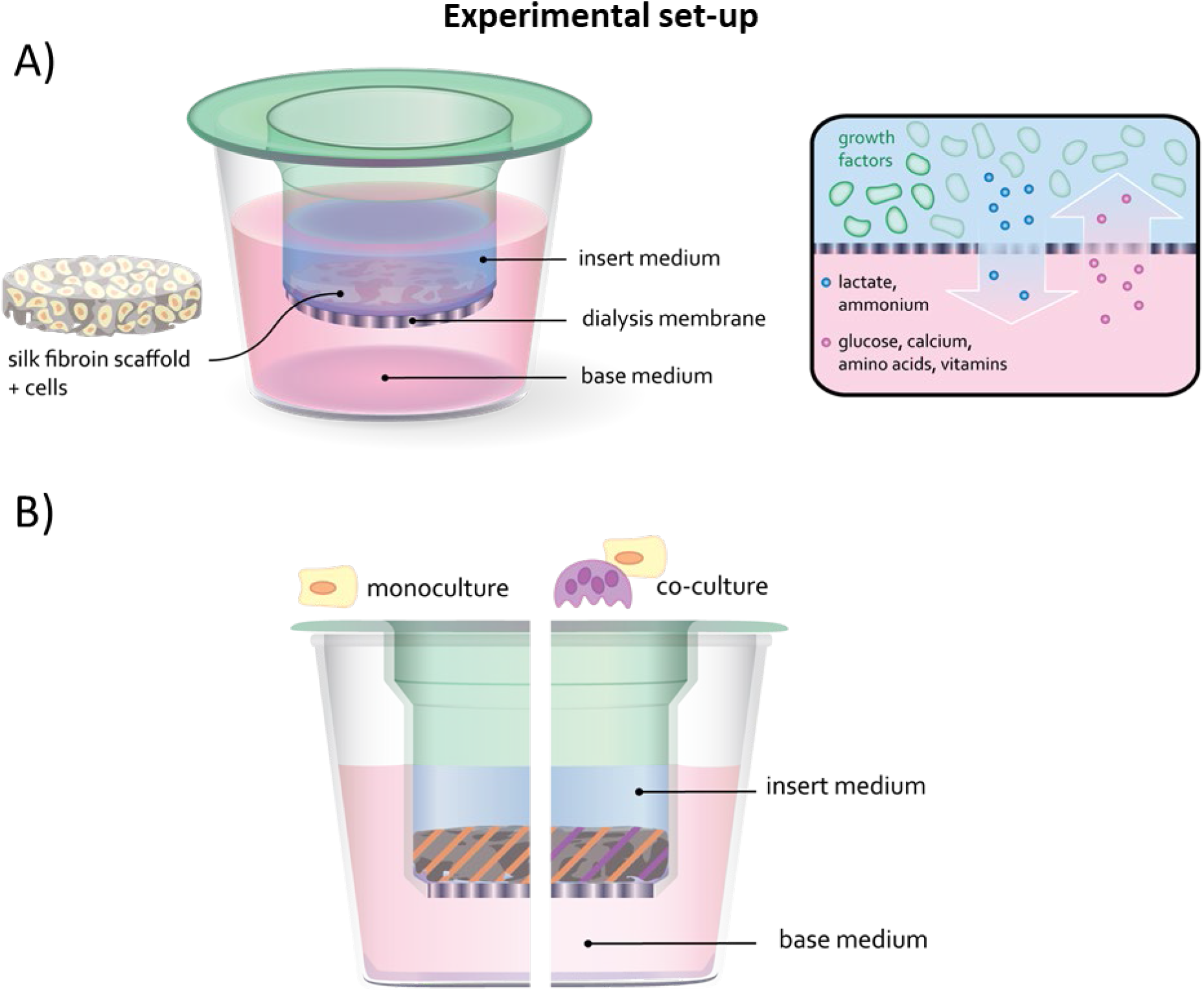
Experimental set-up: A) A silk fibroin scaffold was seeded with cells and cultured in a custom-made dialysis culture dish. The dialysis membrane ensured retaining of high molecular weight growth- and communication factors in the cell culture insert. Nutrients and waste products of low molecular weight can pass the dialysis membrane and will be supplied and removed by diffusion across the membrane. B) In the monoculture experiment the silk fibroin scaffold is seeded with MSCs. In the coculture experiment the scaffold is seeded with MSCs and MCs.

For *in vitro* bone tissue formation, the extracellular matrix (ECM) production by the cells is of uttermost importance. Osteoblasts produce collagen type 1 ECM that is mineralized both intrafibrillarly and extrafibrillarly [12]. The cells’ secretome is essential in this process. Continuous removal or periodic exchange of culture medium considerably disturbs these mineralization processes, making it difficult to induce *in vitro* bone tissue formation [7]. Therefore, in this study, a dialysis culture approach is investigated for an *in vitro* bone remodeling model by differentiation of human mesenchymal stromal cells (MSCs) into osteoblasts and monocytes (MCs) into osteoclasts (Figure 1B). A custom-made simple dialysis culture system with a commercially available cellulose dialysis insert is developed. It is believed that the dialysis of culture medium could contribute to a more efficient and more physiological environment for cell proliferation and differentiation, as the cells’ secretome remains in the culture.

## 2. Materials and Methods

### 2.1 Fabrication of the silk fibroin scaffolds

Silk fibroin scaffolds were produced as previously described [13], [14]. Briefly, Bombyx mori L. silkworm cocoons were degummed by boiling in 0.2M Na_2_CO_3_ for 1 hour. Dried silk was dissolved in 9M LiBr and dialyzed against ultra-pure water (UPW) using SnakeSkin Dialysis Tubing (MWCO: 3.5 kDa; Thermo Fisher Scientific, Breda, The Netherlands). Dialyzed silk solution was snap frozen in liquid nitrogen (−190 °C), lyophilized, and dissolved in 1,1,1,3,3,3-Hexafluoro-2-propanol (HFIP, FCB125463, FluoroChem), resulting in a 17% (w/v) solution. One milliliter silk-HFIP solution was added to 2.5 g NaCl with a granule size between 250 and 300 μm in a Teflon container and allowed to air dry. Silk-salt blocks were immersed in 90% methanol in UPW for 30 minutes to induce β-sheet formation [15]. NaCl was extracted in UPW for 2 days. Scaffolds were cut into disks of 1 mm height, punched with a 5-mm diameter biopsy punch, and autoclaved in phosphate buffered saline (PBS) at 121°C for 20 min for sterilization.

### 2.2 Fabrication of the dialysis culture plates

A holder with 48 holes was custom-made (poly carbonate, designed in Inventor Professional, Autodesk) to fit a 48-wells plate (677180, Greiner) (Figure 2). The holder was autoclaved at 121°C for 20 min and placed on top of a 48-wells plate in a biosafety cabinet. Slide-a-lyzer mini dialysis devices (0.1 mL, 69590, Thermo Fisher scientific) with a MWCO of 20 kDa were sterilized using UV-light on both sides for 15 minutes each. The mini dialysis devices were carefully placed in the holder. All membranes were pre-wetted using sterile PBS which was pipetted in both the base and the insert of the wells (Figure 1). Later, the PBS was removed and replaced with the appropriate cell culture medium.

**Figure 2.**
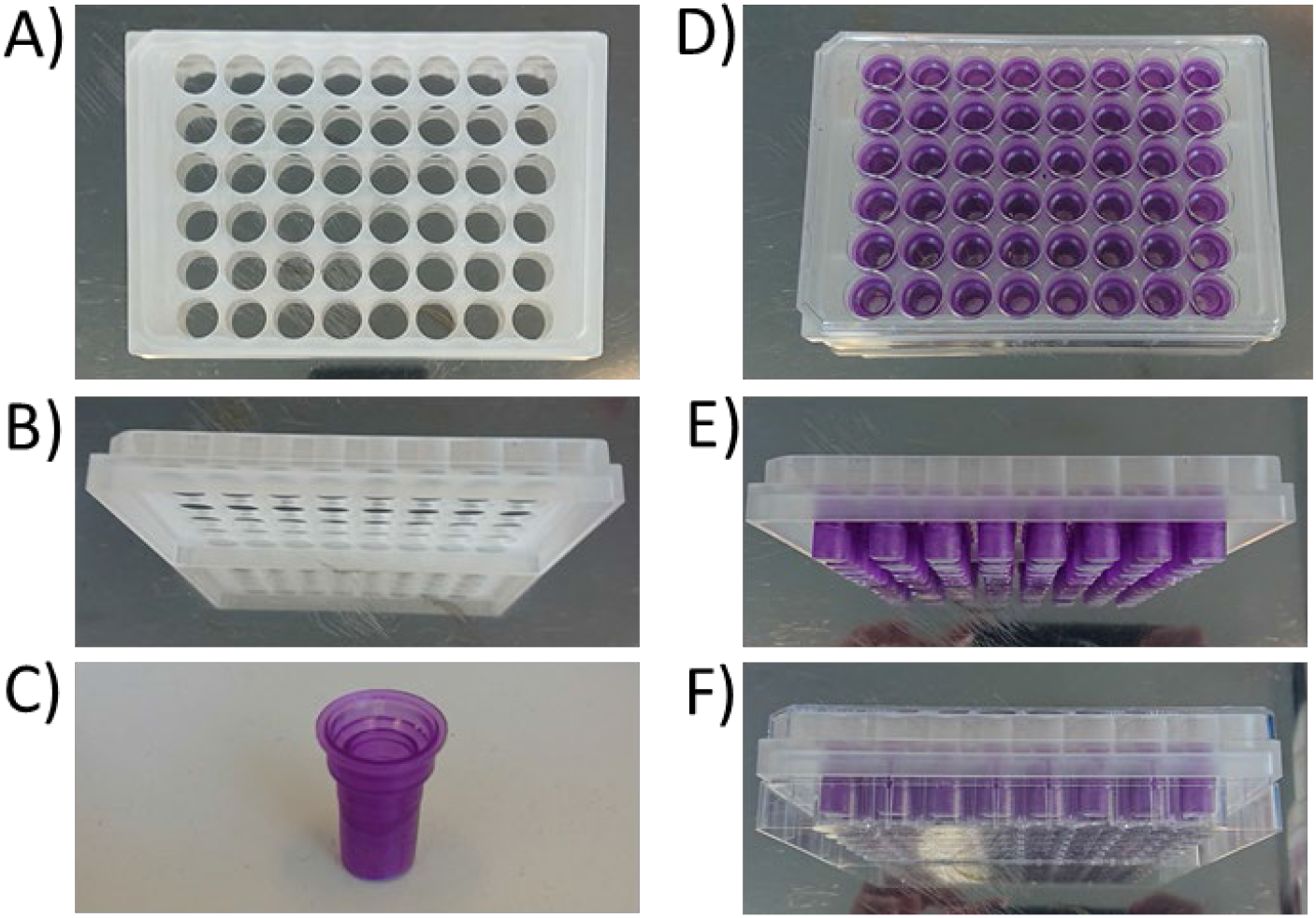
The custom-made holder for the 48 wells plate. A) top view, B) side view, C) dialysis cup, D) top view of holder filled with dialysis cups, E) side view of holder filled with dialysis cups, F) holder with cups in 48 wells plate.

### 2.3 Monoculture: isolation, expansion and cultivation of MSCs

MSC isolation and characterization from human bone marrow (Lonza, Walkersville, MD, USA) was performed as previously described [16]. MSCs were frozen at passage 3 with 1.25*10^6^ cells/ml in freezing medium containing FBS (BCBV7611, Sigma-Aldrich) with 10% dimethylsulfoxide (DMSO, 1.02952.1000, VWR, Radnor, PA, USA) and stored in liquid nitrogen until further use. Before experiments, MSCs were thawed and seeded at a density of 2.5*10^3^ cells/cm^2^ in expansion medium containing DMEM (high glucose, 41966, Thermo Fisher Scientific), 10% FBS (BCBV7611, Sigma Aldrich), 1% Antibiotic Antimycotic (anti-anti, 15240, Thermo Fisher Scientific), 1% Non-Essential Amino Acids (11140, Thermo Fisher Scientific), and 1 ng/mL basic fibroblastic growth factor (bFGF, 100-18B, PeproTech, London, UK) at 37 ºC and 5% CO_2_. After 9 days, cells were detached using 0.25% trypsin-EDTA (25200, Thermo Fisher Scientific) and directly used for experiments at passage 4. A dynamic seeding process was used as previously described [17]. Briefly, SF scaffolds were incubated with a cell suspension (3×10^5^ cells/4 mL expansion medium) in 50-mL tubes placed on an orbital shaker at 150 rpm for 6 hours in an incubator at 37°C and 5% CO_2_. Next, the samples were transferred to the dialysis wells plates and incubated at 37°C and 5% CO_2_ for a total 4 weeks in osteogenic monoculture medium containing DMEM (low glucose, 22320, Thermo Scientific), 10% human platelet lysate [11] (hPL, PE20612, PL BioScience, Aachen, Germany), 1% Anti-Anti, 0.1 μM dexamethasone (D4902, Sigma-Aldrich), 0.05 mM ascorbic acid-2-phosphate (A8960, Sigma-Aldrich), 10 mM β-glycerophosphate (G9422, Sigma-Aldrich). The insert of the dialysis wells plates contained the scaffold and 200 μL osteogenic medium, the base contained 500 μL osteogenic medium. Medium was changed according to Table 1. The culture was maintained for 28 days.

**Table 1.**
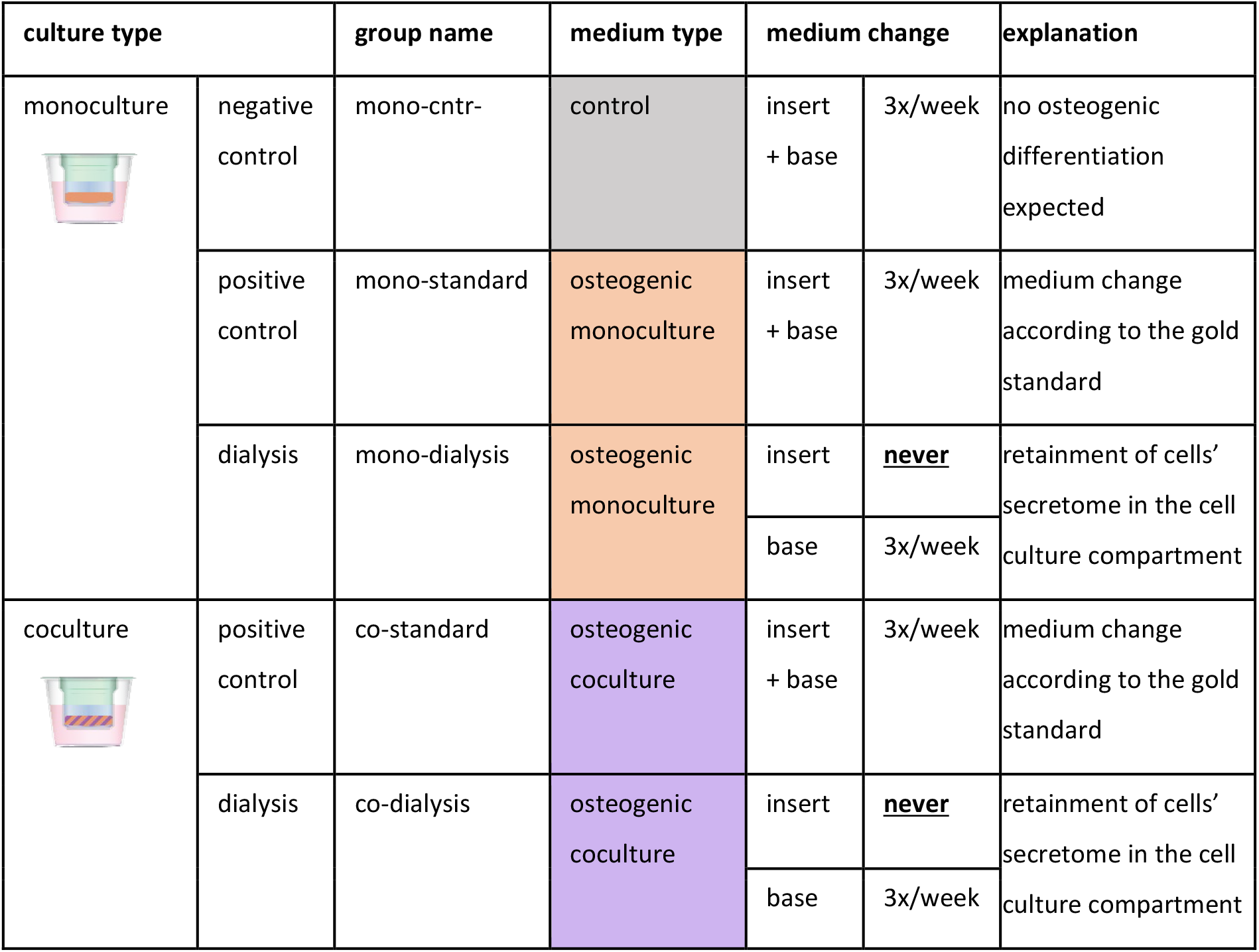
Overview of group name, culture medium type, medium change schedule and explanation

### 2.4 Coculture: isolation of monocytes and cultivation of MSCs + monocytes

Human peripheral blood buffy coats from healthy volunteers under informed consent were obtained from the local blood donation center (agreement NVT0320.03, Sanquin, Eindhoven, the Netherlands). The buffy coats (~ 50 mL) were diluted to 200 mL in 0.6 % (w/v) sodium citrate in PBS adjusted to pH 7.2 at 4 °C (citrate-PBS), after which the peripheral mononuclear cell fraction was isolated by carefully layering 25 mL diluted buffy coat onto 13 mL Lymphoprep (07851, StemCell technologies, Cologne, Germany) in separate 50 mL centrifugal tubes, and centrifuging for 20 min with lowest brake and acceleration at 800×g at RT. Human peripheral blood mononuclear cells (PBMCs) were collected, resuspended in citrate-PBS and washed 4 times in citrate-PBS supplemented with 0.01% bovine serum albumin (BSA, 10735086001, Sigma-Aldrich, Zwijndrecht, The Netherlands) to remove all Lymphoprep. PBMCs were frozen in liquid nitrogen in freezing medium containing RPMI-1640 (RPMI, A10491, Thermo Fisher Scientific), 20% FBS (BCBV7611, Sigma-Aldrich) and 10% DMSO and stored in liquid nitrogen until further use. Prior to experiments, monocytes (MCs) were isolated from PBMCs using manual magnetic activated cell separation (MACS). PBMCs were thawed, collected in medium containing RPMI, 10% FBS (BCBV7611, Sigma-Aldrich) and 1% penicillin-streptomycin (p/s, 15070063, Thermo Fisher Scientific), and after centrifugation resuspended in isolation buffer (0.5% w/v BSA in 2mM EDTA-PBS). The Pan Monocyte Isolation Kit (130-096-537, Miltenyi Biotec, Leiden, The Netherlands) and LS columns (130-042-401, Miltenyi Biotec) were used according to the manufacturer’s protocol. After magnetic separation, the cells were directly resuspended in osteogenic coculture medium (αMEM 41061, 10% hPL, 1% Anti-Anti with 0.1 μM dexamethasone, 0.05 mM ascorbic acid-2-phosphate, 10 mM β-glycerophosphate) spiked with 50 ng/mL macrophage colony stimulating factor (M-CSF, 300-25, PeproTech). The cells were counted and 1.9 × 10^6^ monocytes were resuspended in 20 μL coculture medium and per scaffold added on top of the scaffolds that had previously been seeded with MSCs as described above (chapter 2.4). After 1.5 hour incubation at 37 °C and 5% CO_2_, the rest of the 180 uL medium was added to the insert. 500 uL medium was added to the base. Medium was changed according to Table 1. After 2 days, 50 ng/mL receptor activator of NFκ-B ligand (RANKL, 310-01, PeproTech) was added to the coculture medium and maintained for the rest of the culture. The culture was maintained for 28 days.

### 2.5 Glucose Assay

A glucose assay was used to measure the glucose concentration in the medium at day 7, 14, 21 and 28 in both the insert and the base (n=3). The glucose concentration was measured to ensure passage of glucose as a nutrient component over the dialysis membrane during the 28 day culture period. The method was adapted from Hulme *et al*. [18]. Briefly, on the day of assay, a buffer/chromophore reagent was prepared by mixing an equal volume (3.5 mL) of 4-aminoantipyrine (10 mM, 06800, Sigma–Aldrich) and N-ethyl-N-sulfopropyl-m-toludine (10 mM, E8506, Sigma–Aldrich) with 3.0 mL of 0.8 M sodium phosphate buffer at pH 6.0. 100 μL of this reagent was added to 10 μL horseradish peroxidase (1.6 units/mL, 77332, Sigma-Aldrich) and 2 μL of sample/standard (0–1.25 mM D-glucose (15023-21, Thermo Fisher) in wells of a 96-well plate. The reaction was initiated by the addition of 10 μL glucose oxidase (2.7 units/mL, G7141-10KU, Sigma-Aldrich) and after 30 min incubation at room temperature in the dark, absorbance was measured at 550 nm using a plate reader (Synergy HTX, Biotek). Glucose concentration of each sample was then calculated using a standard curve.

### 2.6 Lactate Assay

A lactate assay was used to measure the lactate concentration in the medium at day 7, 14, 21 and 28 in both the insert and the base (n=3). The lactate concentration was measured to ensure passage of lactate as a metabolic waste product over the dialysis membrane during the 28 day culture period. The methods was adapted from Salvatierra *et al*. [19]. Briefly, a reaction mix was prepared containing 5 mg/mL of β-Nicotineamide Adenine Dinucleotide (N7004,1G, Sigma-Aldrich), 0.2 M glycine buffer (G5418, Sigma-Aldrich), and 22.25 units/mL of L-Lactic Dehydrogenase (L3916, Sigma-Aldrich). A standard curve was prepared using 20 mM lactate stock solution by adding sodium L-lactate (L7022, Sigma-Aldrich) to dH_2_O. Equal parts (40 μL) of the reaction mix and each sample/standard were mixed in a 96-well plate. After 30 minutes of incubation at 37°C, absorbance was measured at 340 nm using a plate reader (Synergy HTX, Biotek). Lactate concentration of each sample was then calculated using a standard curve.

### 2.7 Cell metabolic activity

PrestoBlue assay was used to analyze and track the metabolic activity of the MSCs and MCs in the SF scaffold. Briefly, at days 7, 14, 21 and 28, samples (n=4 per group) were transferred to a clean 48 wells plate filled with 200 μL of 10 v/v % PrestoBlue reagent (A13262, Thermo Fisher) in the appropriate medium (mono or coculture) and incubated for 1 h at 37 °C. In a 96-well assay plate 100 μL of the PrestoBlue solution was added in duplo per sample and the absorbance was measured at 570 and 600 nm. Dye reduction rate was calculated according to manufacturer’s instructions.

### 2.8 Cell-associated alkaline phosphatase activity

Alkaline phosphatase (ALP) was used to indirectly quantifying osteoblast activity. At days 7, 14, 21 and 28, after the Presto Blue assay (see 2.7 Cell metabolic activity) scaffolds (n=4 per group) were washed in PBS and disintegrated in 0.5 mL of 0.2% (v/v) Triton X-100 and 5 mM MgCl_2_ solution using steel beads and a Mini Beadbeater™ (Biospec). The remaining solution was centrifuged at 3000g for 10 minutes. In a 96-well assay plate, 80 μL of the supernatant was mixed with 20 μL of 0.75M 2-amino-2-methyl-1-propanol buffer (A65182, Sigma-Aldrich) and 100 μL 100 mM p-nitrophenylphosphate solution (71768, Sigma-Aldrich) and incubated for 10 minutes, before adding 100 μL 0.2M NaOH stop solution. Absorbance was measured at 405 nm using a plate reader (Synergy HTX, Biotek) and these values were converted to ALP activity (converted p-nitrophenyl phosphate in μmol/ml/min) using standard curve absorbance values.

### 2.9 DNA assay

The solution obtained for the ALP activity assay (see 2.8) after disintegration of the scaffold was further used the measure the amount of DNA as an attribute for cell number. In each Eppendorf tube, 340 uL of liquid remained after the ALP measurements. Next, 340 uL of papain (280 ug/mL, p4762, Sigma-Aldrich) in digestion buffer was added and the samples were incubated overnight at 60 °C under constant shaking in a shaking Eppendorf tubes water bath. The DNA content of the supernatant was determined using the Qubit HS dsDNA Assay Kit (Q32854, Invitrogen, Thermo Fisher) according to the manufacturer’s instructions. The calculated ALP activity was normalized by the DNA content.

### 2.10 Tartrate-resistant acid phosphatase activity

Tartrate-resistant acid phosphatase (TRAP) was used to indirectly quantifying osteoclast activity. Supernatant medium samples of both the insert and the base were taken at medium change and stored at −80 °C at day 7, 14, 21, and 28. 100 μL pNPP buffer (1 mg/mL p-nitrophenylphosphate, 0.1 mol/L sodium acetate, 0.1 % (v/v) Triton-X-100 in PBS, first adjusted to pH 5.5, then supplemented with 30 μL/mL tartrate solution (Sigma Aldrich)) and 20 μL culture medium or nitrophenol standard in PBS were incubated in 96-well assay plates at 37 °C. After 90 min, 100 μL 0.3 mol/L NaOH was added to stop the reaction. Absorbance was read at 405 nm using a plate reader and absorbance values were converted to TRAP activity (converted p-nitrophenyl phosphate in μg/ml) using standard curve absorbance values.

### 2.11 Immunohistochemistry

At day 28, scaffolds (n=4 per group) were washed with PBS and immersed first in 5% and then in 35% sucrose solution in PBS at room temperature for 10 min each. The scaffolds were embedded in cryomolds containing Tissue-Tek OCT compound (Sakura, The Netherlands), snap frozen in liquid nitrogen, cut into 10 μm thick sections using a Cryotome Cryostat (Leica biosystems), and mounted on Superfrost Plus microscope slides (Thermo Fisher Scientific). Next, sections were immunostained after washing with PBS-tween, fixing in 10% neutral-buffered formalin for 10 min at room temperature, washing again with PBS-tween, permeabilizing in 0.5% Triton X-100 in PBS for 10 min and blocking in 10% normal goat serum in PBS for 30 min. Cells were incubated with DAPI, Phalloidin and immunostainings (Table 2) in PBS for 1 h. Images were taken with either a Zeiss Axio Observer 7 (collagen type-1 images) or a Leica TCS SP5X (RUNX-2 and osteopontin images).

**Table 2.** List of all dyes and antibodies used

| Antigen | Source | Catalogue No | Label | Species | Concentration/Dilution |
| --- | --- | --- | --- | --- | --- |
| DAPI | Sigma-Aldrich | D9542 |  |  | 0.1 µg/mL |
| Atto 647<br>conjugated<br>Phalloidin | Sigma-Aldrich | 65906 |  |  | 50 pmol |
| Collagen<br>type-1 | Abcam | Ab34710 |  | Rabbit | 1:200 |
| RUNX-2 | Abcam | Ab23981 |  | Rabbit | 1:500 |
| Osteopontin | Thermo<br>Fisher | 14-9096-82 |  | Mouse | 1:200 |
| Anti-rabbit<br>(H+L) | Molecular<br>Probes | A11008 | Alexa 488 | Goat | 1:200 |
| Anti-mouse<br>IgG1 (H+L) | Molecular<br>Probes | A21127 | Alexa 555 | Goat | 1:200 |

### 2.12 Histology

Cryosections made as described (2.11 Immunohistochemistry) were washed with PBS, fixed in 10% neutral-buffered formalin for 10 min at room temperature, washed again with PBS, and stained with Alizarin Red (2% in distilled water, A5533, Sigma-Aldrich) for 15 minutes to identify mineralization. Sections were imaged using a Zeiss Axio Observer Z1 microscope.

### 2.13 Scanning electron microscopy

Constructs for scanning electron microscopy (SEM) (n=4 per group for cocultures) were fixed at day 14 in 2.5 % glutaraldehyde for 24 h at 4 °C, dehydrated with a graded ethanol series (2 × 50 %, 70 % and 95 %, 3 × 100 % 10-15 min each) followed by a graded 1,1,1-Trimethyl-N-(trimethylsilyl)silanamine (HMDS)/ethanol series (1 : 2, 1 : 1, 2 : 1, 3 × 100 % HMDS 15 min each), dried at room temperature overnight and sputter coated with 5 nm gold (Q300TD, Quorum Technologies Ltd, Laughton, UK) prior to imaging with SEM (Quanta600, FEI Company, Eindhoven, the Netherlands) with a spot size of 3.0, 10.00 kV, working distance 10 mm. Images were colored using Adobe Photoshop 2022.

### 2.14 Statistical analysis

Statistical analyses were performed, and graphs were prepared in GraphPad Prism (version 9.4.0, GraphPad, La Jolla, CA, USA) and R (version 4.0.2). Data were tested for normality in distributions with Shapiro-Wilk tests and for equal variances with Levene’s tests. Glucose, lactate (Figure 3) and TRAP data (Figure 6) were normally distributed without equal variances and differences were therefore tested with Welch’s t-tests and presented with mean and standard deviation. Cell metabolic activity data (Figure 4) was not normally distributed and without equal variances and differences were tested with Mann-Whitney tests and presented with median and interquartile range. ALP (Figure 5) data was normally distributed and variances were equal and differences were tested with two-way ANOVA and Tukey post hoc test for multiple comparisons and presented as mean and standard deviation.

**Figure 3.**
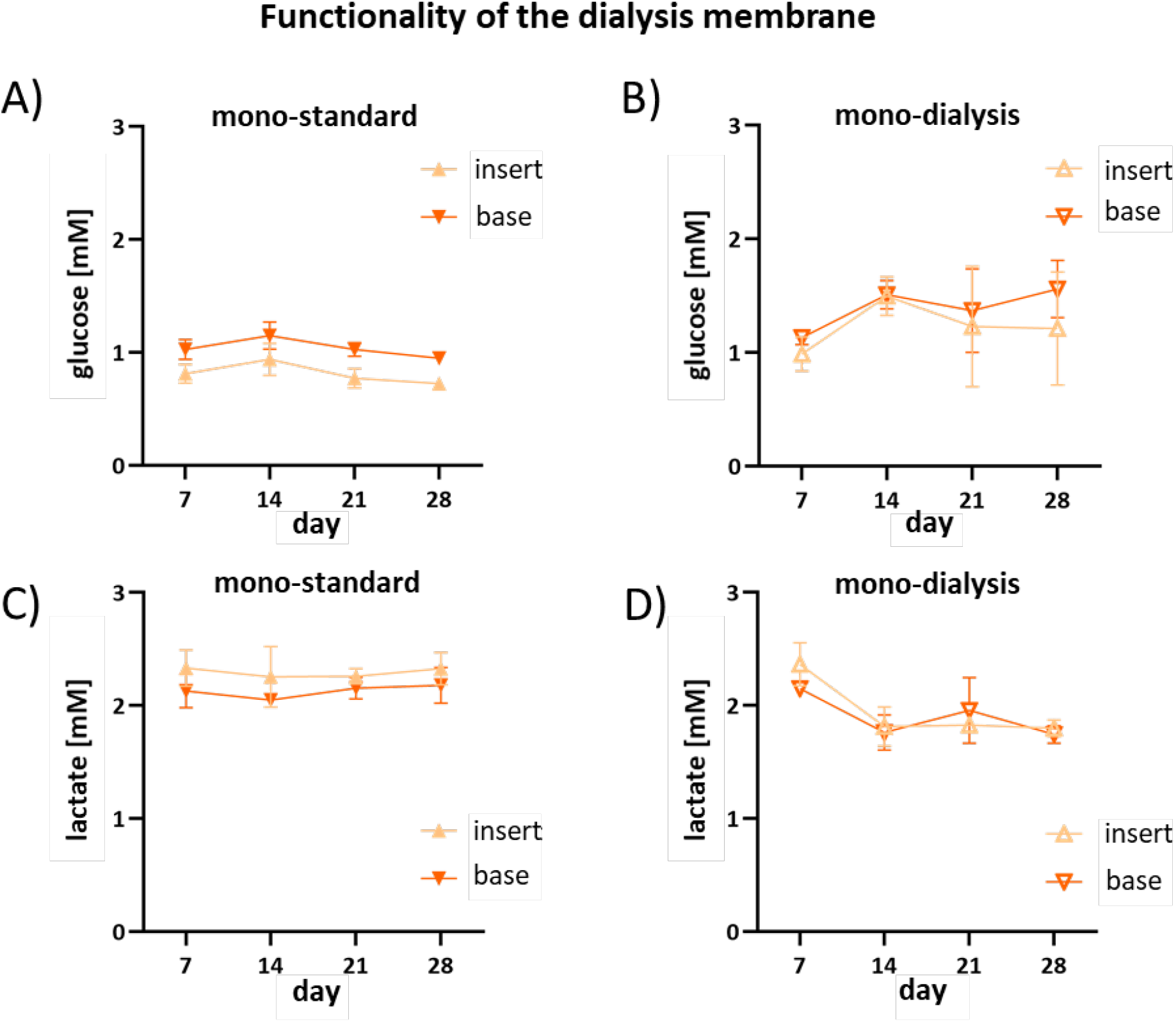
Functionality of the dialysis membrane A) + B) Glucose and C) + D) lactate measurements on culture medium of the monoculture cell experiment over 28 days performed in our dialysis system (n=3). Measurements were taken in both the insert and the base of the system. At each timepoint the difference between insert and base is non-significant (p>0.05, Welch’s t-tests).

**Figure 4.**
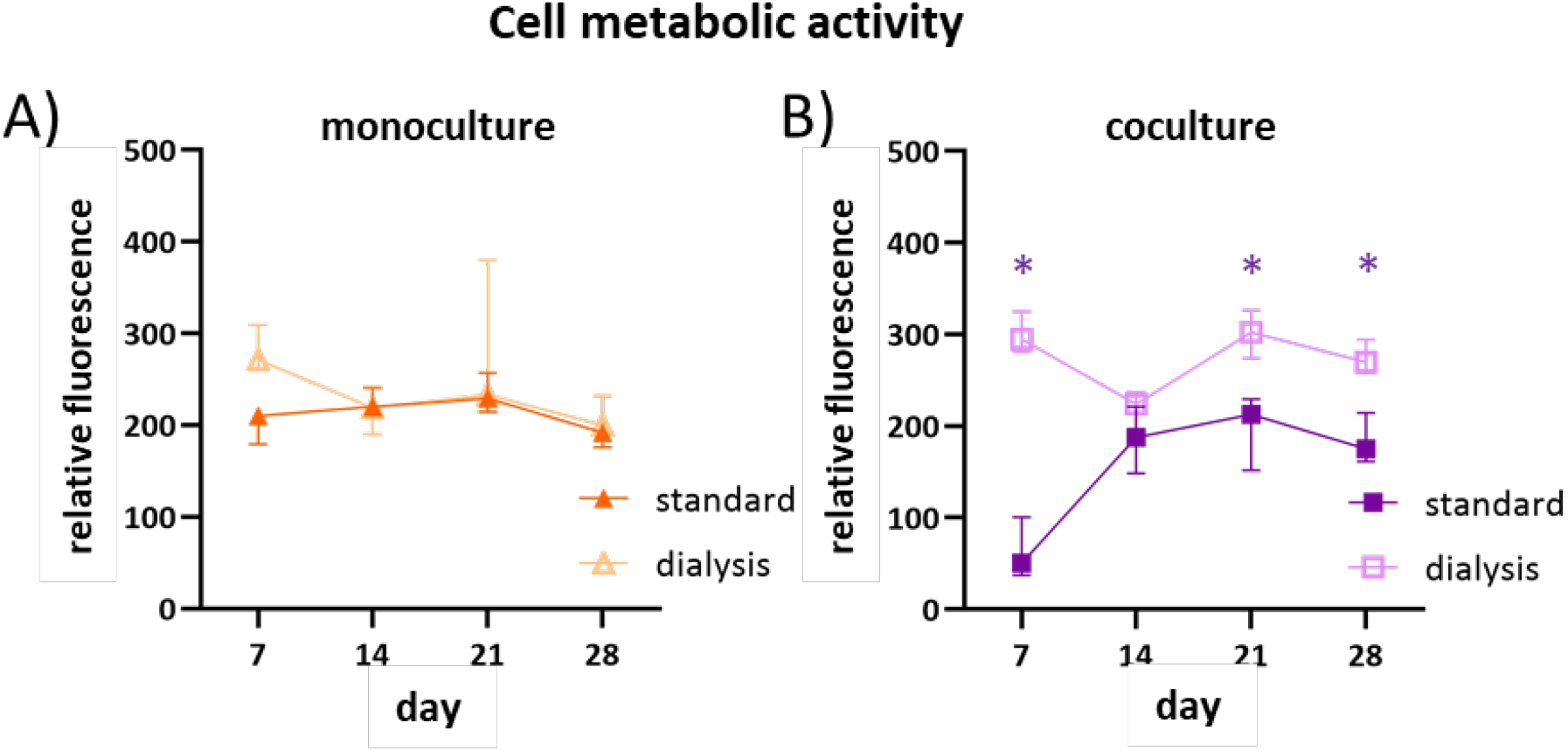
Cell metabolic activity (presto blue) of A) the monoculture and B) coculture measured during 28-day cell experiments shown as relative fluorescence vs. medium control (n=4). At each timepoint the difference between the control and dialysis samples was determined, with significant differences for p<0.5 (Mann-Whitney tests) indicated with an asterisk.

**Figure 5.**
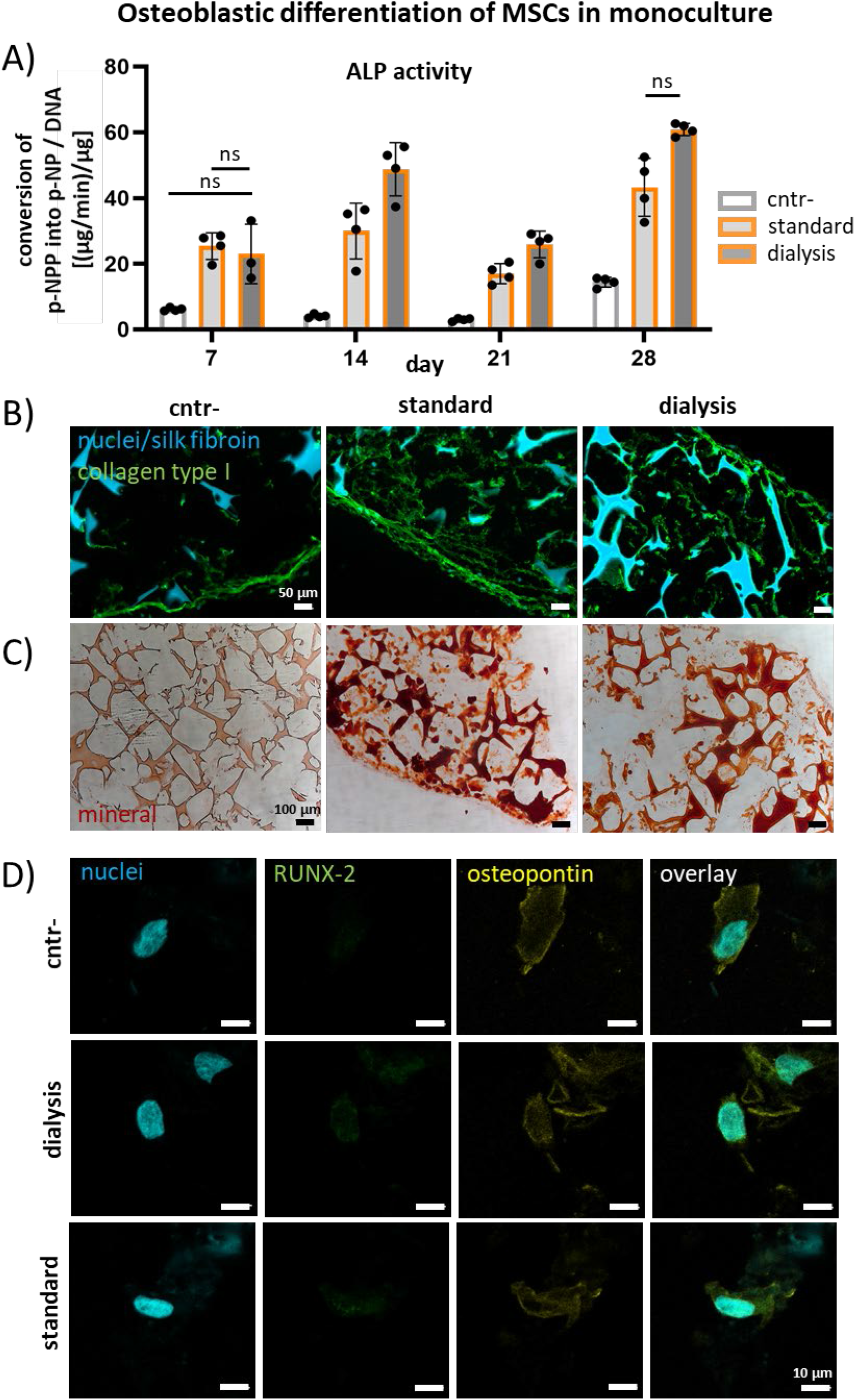
Osteoblastic differentiation of MSCs in monoculture. A) ALP activity divided by DNA (n=4); significant differences are determined by two-way ANOVA with Tukey’s post hoc test. All differences are significant (p<0,05), unless indicated with “ns”. B) Collagen type 1 production by the cells visualized by immunostaining at day 28. C) Mineralization shown by Alizarin Red staining at day 28. D) RUNX-2 and osteopontin expression visualized by immunostaining at day 28.

**Figure 6.**
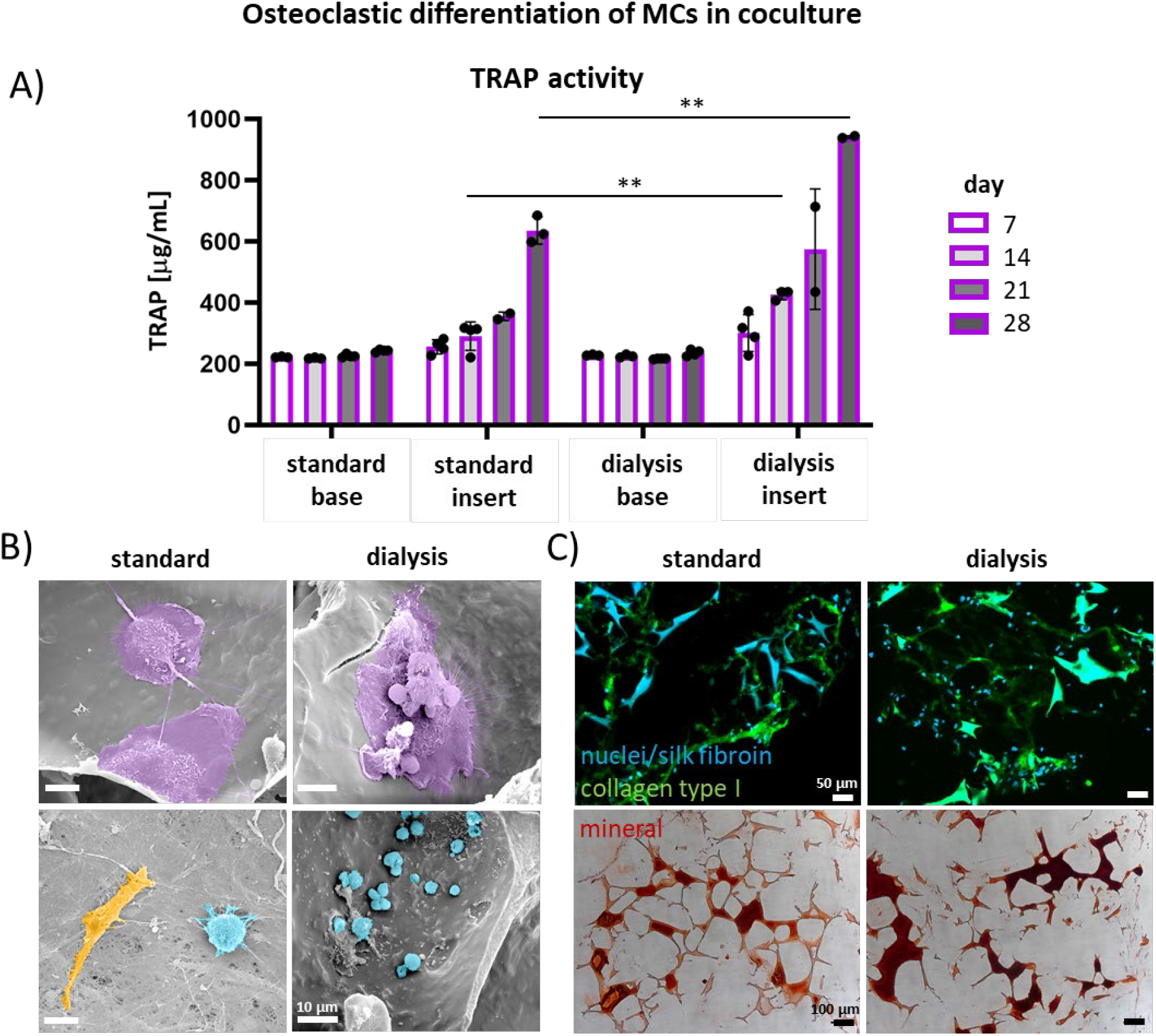
Osteoclastic differentiation of MCs in coculture. A) TRAP activity; significant differences (p<0.05) are determined by comparing the inserts the of the standard to the dialysis group for each timepoint using Welch’s tests. B) Collagen type 1 production of the cells visualized immunostaining at day 28. C) Mineralization shown by Alizarin Red staining at day 28. D) Cell morphology visualized by SEM at day 14. Osteoclast-like cells are depicted in purple, osteoblast-like in orange and monocyte-like in blue.

## 3. Results

### 3.1 Functionality of the dialysis membrane: continuous passage of glucose and lactate

To test whether the dialysis membrane did not clog over time, glucose and lactate passage over the membrane were measured over the culture period of 28 days in the monoculture. Measurements were taken from the cell culture media of both the insert and the base of the dialysis system (Figure 1). Free passage of glucose and lactate molecules over the membrane was indicated by non-significant differences of the measured amount of glucose and lactate between the insert and the base medium in all groups (Figure 3A-D).

### 3.2 Functionality of the dialysis membrane: cell metabolic activity

To ensure that no toxic components build up in the cell culture system, cell metabolic activity was followed over the culture time. Cell metabolic activity was measured in both the mono- and coculture for the standard and dialysis groups. In the monoculture, the cell metabolic activity of the standard group compared to the dialysis group was similar for each timepoint with no significant differences (Figure 4A). In the coculture, dialysis seems to be beneficial compared to the standard as the cell metabolic activity of the dialysis group was significantly higher (co-standard vs. co-dialysis) on day 7, 21, 28 (Figure 4B). The results indicate that both in mono- and coculture, the cells were equally or at some timepoints more metabolically active in the dialysis group compared to the standard showing that the dialysis system could beneficially influence the cell metabolic activity.

### 3.2 Osteoblastic differentiation of MSCs in monoculture

ALP activity is a widely used marker for indirectly quantifying osteoblastic activity [20]. ALP activity on the cells’ surface was determined weekly in the monoculture over a 28 day time period and normalized against the amount of DNA. The ALP activity was lowest in the negative control throughout all timepoints and was only slightly elevated at day 28. As expected, the ALP activity in the standard and dialysis groups was increased and this difference was significant compared to the negative control starting from day 14. There was a trend that the ALP activity was slightly higher in the dialysis group compared to the standard group with significant differences at day 14 and 21 (Figure 5A). These results indicate higher osteoblastic activity in the dialysis group compared to the standard group (Figure 5A).

To confirm extracellular matrix production and osteoblastic differentiation of the MSCs in the monoculture, samples were stained for collagen type-1, mineralization (Alizarin Red staining), RUNX-2 and osteopontin expression at the endpoint of the experiment (day 28). Collagen type 1 was present in all groups, but seemingly less and mostly located at the outer rim of the scaffold in the negative control compared to the standard and dialysis groups (Figure 5B). Mineralization was visible in the standard and dialysis groups while being absent in the negative control (Figure 5C). RUNX-2 expression was faintly present in the nucleus of the standard and dialysis groups while being absent in the negative control (Figure 5D). Osteopontin expression was visible in the body of cells in all groups (Figure 5D). The results confirm that the presence of the dialysis system allows for MSC to produce an ECM composed of collagen and mineral and for the cells to differentiate into osteoblasts.

### 3.3 Osteoclastic differentiation of MCs in coculture

TRAP activity is a widely used marker for indirectly quantifying osteoclastic activity [20]. TRAP activity was measured weekly in the coculture medium of both the base and the insert over a 28 day time period. Over time, TRAP activity in the base of both groups was stable and low (Figure 6A). In the insert TRAP activity increased over time with a significantly higher activity in the dialysis group compared to the standard at day 14 and 28 (Figure 6A). These results suggest retainment of TRAP in the insert of the dialysis group leading to higher osteoclastic activity compared to the standard group, where the medium in the insert was replaced at every medium change (Figure 6A).

The coculture was investigated with SEM at day 14. In both the standard and dialysis group large (~25-40 μm) cells were visible that were attached to the surface (Figure 6B, in purple). These cells resemble osteoclast in morphology and size. Also, smaller (~5-10 μm) rounded cells were visible in both groups. These cells resemble monocytes in morphology and size and were expected to be present as monocytes had been seeded in abundance (Figure 6B, in blue). Further, an elongated cell was seen, resembling a towards osteoblast differentiated MSC in morphology and size (Figure 6B, in orange).

For the coculture, a different type of culture medium was used compared to the monoculture, as MSCs and MCs need different components to differentiate into osteoblasts and osteoclasts respectively. To confirm extracellular matrix production of the MSCs in the coculture medium, samples were stained for collagen type 1 and mineralization (Alizarin Red staining) at the endpoint of the experiment (day 28). In both coculture conditions, the standard and the dialysis, an ECM was produced with collagen type 1 and mineralization (Figure 6C).

## 4. Discussion

Regular culture medium exchange leads to loss of valuable auto- and paracrine factors produced by the cells. However, frequent renewal of culture medium is necessary to prevent waste product accumulation and to supply fresh nutrients and is therefore the gold standard in cell culture applications. The use of a cell culture compartment with a dialysis membrane could overcome the need for frequent cell culture medium renewal in the cell culture compartment as low MW molecules such as nutrients and waste products can easily pass the membrane, while high MW components such as growth factors are maintained. Here, a coculture system of human osteoblasts and osteoclasts was established in a dialysis system that seemed to allow for maintaining of the cells’ secretome. Typical components of the ECM together with typical enzymatic activity of the cells (ALP and TRAP activity) were shown.

In the present study, a simple cell culture system with a dialysis membrane was developed. For the use of a dialysis membrane in long-term cell culture applications, the characteristics of the membrane are important. Based on the sizes of the osteogenic cell culture medium components, a membrane with a MWCO of 20 kDa was chosen. The supplements of osteogenic monoculture medium are assumed to be able to pass this membrane: dexamethasone (390 Da), ascorbic acid (176 Da) and β-glycerophosphate (172 Da). However, the coculture supplements RANKL (20 kDa) and MCS-F (37 kDa) are not and are therefore only added in the beginning of the culture. No literature was found about the long-term half-life of these factors at 37°C, only that no degradation was detected within the first 48 hours [21], [22]. Although RANKL and MCS-F were only added in the beginning, evidence for the presence of osteoclasts was found. Thus, the supplements either stayed active or stimulated the coculture enough in the beginning. The possible cells’ secretome consists of high MW molecules that are assumed not to be able to pass the membrane such as ALP (86 kDa), RUNX-2 (57 kDa), osteopontin (55 kDa), TRAP (30-35 kDa) and extracellular vesicles (large range of MW).

Dialysis membranes are designed to be non-fouling. However, cell culture medium consists of nutritious liquids which are slightly viscous and sticky and could cause the membrane to clog over time [23]. Also, cell mediated mineral deposition on the cell side of the dialysis membrane has been reported [24]. However, it was not reported whether this deposition influenced the function of the dialysis membrane. Our system was functional over the culture period of 28 days. Small molecules such as glucose (180 Da) and lactate (90 Da) were able to pass the membrane continuously over the whole culture time. Large molecules such as TRAP (30-35 kDa) were maintained in the cell culture compartment. Moreover, the cells remained metabolically active, an indicator that the membrane still provided nutrients and removed waste products.

The dialysis culture system was used to study osteogenic differentiation of MSCs and MCs into osteoblasts and osteoclasts. It was hypothesized that the maintenance of auto- and paracrine factors in the culture medium would contribute to a more physiological microenvironment for the cells. The data reported here revealed higher osteoblastic and osteoclastic activity in the dialysis groups compared to the standard, shown by significantly higher ALP and TRAP activity respectively. Therefore, the dialysis system contributes to an excellent cell culture environment. Remarkably, this significant effect seen in the biochemical assays was not visible in the matrix production (collagen type 1 and mineralization). We hypothesize that this might be due to a difference in overall concentration of ascorbic acid and β-glycerophosphate. In the dialysis system, only the base medium is exchanged (500 μL) while in the standard system both the insert and base media are exchanged (200+500 μL), resulting in a lower overall concentration of these to factors. Ascorbic acid has been shown to increase the secretion of collagen type 1 and β-glycerophosphate is the phosphate source that is needed for mineralization [25]. It is recommended to study this effect in future experiments.

To analyze *in* vitro models over time, non-destructive methods including medium analysis are desired [26]. A limitation of the current dialysis system is that the possibility to take medium samples from the insert is very limited. The principle of the system relies on the fact that the insert stays undisturbed. Also, the current design allows for only 200 μL of cell culture medium in the insert. Taking medium samples would lead to a necessity for adding fresh medium. Therefore, at each timepoint samples were sacrificed to enable the performance of assays on the medium. To overcome this issue, non-invasive assays or sensors could be used such as for example biosensing by particle mobility (BPM) [27] or aptamer based sensors [28]. Such sensors would allow for continuous monitoring of specific biomarkers without the need for medium sampling. A simpler solution could be the use of a larger volume culture medium in the insert. The amount of medium sampled should be very small relative to the total amount of medium in the insert, so that it would not lead to disturbance of the microenvironment. However, making the volume too large, could lead to loss of the effect of the cells’ secretome on the microenvironment due to dilution [1].

A crucial process of *in vitro* bone tissue formation is ECM production and mineralization. The data presented in this study, mainly in the coculture, indicate limited mineralization (alizarin red) in both the standard as the dialysis system. Furthermore, this mineralization occurs primarily in the scaffold material and not in the matrix produced by the cells. There are two possible explanations for this observation. Firstly, the presented experimental set-up cultures statically. It has been shown that for mineralized matrix production the cells prefer mechanical stimulation [29]. Here, the MSCs were initially stimulated by dynamic seeding [17], but this effect was limited compared to what is usually seen in a dynamically loaded environment [30]. In future experiments, the use of a bioreactor that can apply for example fluid flow induced shear stress would be desired to improve the mineralization [30]. But implementing the dialysis system with its membrane will concomitantly affect the fluid flow. Secondly, it is still a major challenge to use the right type of coculture medium as both the MSCs and the MCs should be stimulated to differentiate into osteoblasts and osteoclasts respectively. The coculture medium used may not have offered the ideal balance yet, probably resulting in a less mineralized matrix compared to the monoculture. MCs need a mineralized surface to attach to and to become osteoclasts. Recently, de Wildt *et al*. reported a pre-mineralized SF scaffold that acts as a bone-mimetic template [31]. They used poly aspartic acid and simulated body fluid to pre-mineralize silk fibroin scaffolds before cell seeding. The mineralized scaffolds supported both osteoclastic resorption and osteoblastic mineralization while in our study, the monocytes relied on MSCs first differentiating into osteoblasts and producing a mineralized matrix. Using the pre-mineralized template in combination with a dialysis culture system could lead to an even faster method to generate a physiological bone remodeling model.

The proposed dialysis culture system is not limited to bone and could be beneficial for a wide variety of cell culture applications. There is a huge variety of existing materials and MWCO options for the membrane, making it customizable for different needs. The MWCO can be chosen based on the cell culture medium ingredients and expected secretome. Caution however has to be taken to possible toxic elements in the manufactured dialysis membranes, such as glycerol, which has been shown to inhibit cell proliferation in several cell lines [32] [33].

In conclusion, we have demonstrated the feasibility of a simple to use dialysis cell culture system for bone tissue engineering applications. The dialysis system enables retention of the cells’ secretome and thereby omits the extra effort that the cells have to make to restore their communication after culture medium exchange. The system creates a stable microenvironment for the cells to differentiate into the osteogenic and osteoclastic lineage. This simple culture system has the potential to be applied in other TE fields and is recommended to be used for differentiation of various cell types.

## 7. Author contributions

MV, BW, KI and SH contributed to conception, methodology and design of the study. MV performed and analyzed the experiments. BW provided expertise on the practical experimental work, particularly the coculture experiments. MV wrote the original draft of the manuscript and prepared the figures. All authors contributed to manuscript revision and approved the submitted version. KI and SH contributed to the supervision. SH acquired funding for this research.

## 8. Acknowledgements

We thank Jurgen Bulsink for the design and fabrication of the 48-well plate holder.

